# Quantifying control circuit regulation in the human brain

**DOI:** 10.1101/2021.03.30.437729

**Authors:** Rajat Kumar, Helmut H. Strey, Lilianne R. Mujica-Parodi

**Affiliations:** Department of Biomedical Engineering, Stony Brook University, Stony Brook, NY—USA; Laufer Center for Physical and Quantitative Biology, Stony Brook University, Stony Brook, NY—USA; Athinoula A. Martinos Center for Biomedical Imaging, Department of Radiology, Harvard Medical School, Massachusetts General Hospital, Charlestown, MA—USA

**Keywords:** fMRI, circuit, control, controllability, dysregulation, perturbation, stress

## Abstract

As a field, control systems engineering has developed quantitative methods to characterize the regulation of systems or processes, whose functioning is ubiquitous within synthetic systems. In this context, a control circuit is objectively “well regulated” when discrepancy between desired and achieved output trajectories is minimized, and “robust” to the degree that it is able to regulate well in response to a wide range of stimuli. Most psychiatric disorders are assumed to reflect dysregulation of brain circuits. Yet, probing circuit regulation requires fundamentally different analytic strategies than the correlations relied upon for analyses of connectivity and their resultant networks. Here, we demonstrate how well-established methods for system identification in control systems engineering may be applied to functional magnetic resonance imaging (fMRI) data to extract generative computational models of human brain circuits. These provide two quantitative measures of direct relevance for psychiatric disorders: a circuit’s sensitivity to external perturbation and its dysregulation.

## Main

Control systems engineering, as a field, has developed quantitative methods to characterize the regulation of systems or processes, whose functioning is ubiquitous within synthetic systems, including thermostats and cruise control systems (where the system is designed to use excitatory and inhibitory inputs from a controller to regulate up or down to a desired set point), as well as complex sensory-motor “cognitive” control processes in robotics. What all of these systems have in common is a controller with feedback that modulates inputs to achieve a desired aim. In this context, a control circuit is objectively “well regulated” when discrepancy (error) between desired and achieved output trajectories is minimized, and “robust” to the degree that it is able to regulate well in response to a wide range of inputs (stimuli, perturbations).

Previous work on network control theory^1-3^ has used optimal control to model network controllability in terms of the energy requirements for shifting one network configuration to another (e.g., mapping easy-to-reach versus difficult-to-reach brain states). This work is highly relevant to understanding the brain’s state transitions in response to exogenous inputs, as the network’s structure identifies optimal “control” nodes, whose stimulation exerts disproportionate influence upon the system dynamics. At the same time, controllability omits key features biomimetic to actual neural circuits. Network controllability^2^ thus far has been defined solely with respect to structural constraints from diffusion tensor imaging and how the network topology affects dynamic trajectories along the energy landscape. Yet circuits in the brain, at all scales, include excitatory and inhibitory influence, lags, amplifiers, gates, filters, as well as positive and negative feedback: anatomically specific control elements whose variation in functioning is not captured by whole-network trajectories.

Extending the capabilities of functional magnetic resonance imaging (fMRI) from connectivity to circuits matters because the assumption is that brain circuit dysregulation plays a key role in the etiology of most psychiatric disorders^4-6^. Indeed, disorders outside the brain often involve a breakdown in the control circuit’s ability to maintain homeostatic control^7^ to a specific setpoint (e.g., glucose levels in diabetes) or allostatic regulation^8^ in the control circuit’s ability to not only respond to the environment but also return efficiently to baseline (e.g., autonomic regulation in heart disease). Two reasons to suspect that at least some psychiatric disorders implicate allostatic disruption^9^ are their vulnerability to perturbation in the form of emotional stress (commonly seen in first-break psychosis and relapse) and trajectories with prominent oscillating features (e.g., bipolar disorder and addiction). Candidate circuits that have been consistently implicated in psychiatrically relevant symptoms include the prefrontal-limbic circuit^10,11^ (threat processing, as per anxiety and paranoia), the meso-corticolimbic circuit^12,13^ (reward processing, as per anhedonia and addiction), the cortico-thalamic loop^14,15^ (sensory processing, as per attentional deficits and hallucinations), and combinations thereof. As evident even from the nomenclature, all three circuits have significant overlap as well as mutual feedback. Regardless of which circuit(s) is chosen for study, however, probing their regulation will require analytic strategies that differ fundamentally from the correlations relied upon for analyses of connectivity and their resultant networks. Ultimately, the value added by neuroimaging-derived control circuits is the promise to predict trajectories, not only to validate the field’s models but also to have them predict individual variability in clinical outcomes.

Here, we demonstrate how well-established methods for system identification^16^ in control systems engineering may be applied to fMRI data. In doing so, we will generate two quantitative measures of direct relevance for psychiatric disorders: the circuit’s sensitivity to external perturbation (its responsivity to stimuli or “stress”), and its control error (how well it achieves its desired state, with respect to both processing information and maintaining allostasis). As shown in **Fig. 1**, our general strategy has three parts. The first step is to identify the circuit architecture. For all nodes, we determine each node’s direct inputs using dynamic causal modeling^17^ (DCM). Thus, we define the basic structure, including feedback loops. The second step is to identify the circuit functionality (in control systems engineering, this process is called “system identification”). We use a linear state-space model to obtain parameters for the difference equation (transfer function) that best specifies how each node transforms its inputs to outputs. This transfer function, which takes the form of an autoregressive model with exogeneous inputs (ARX), describes each node by a finite number of hidden dynamic eigenmodes with characteristic frequencies and damping ratios. These transfer functions link together to form a putative circuit, which we can validate by predicting the system’s response against independent validation datasets. The third step is to probe the system. By testing the system with a gradient of inputs of different frequencies (impedance measurements) and colored noise, we explore the full landscape of the circuit’s possible behaviors. In particular, we want to pay attention to which parts of the control circuit are exponentially growing, which are decaying, and which are oscillating. By calculating the impact of the system’s damping ratio and frequency ratio on the modulation of inputs, we obtain a measure of sensitivity to external perturbation. Finally, we calculate the control error, which provides a quantitative measure of how well regulated or dysregulated the circuit is.

**Fig. 1:**
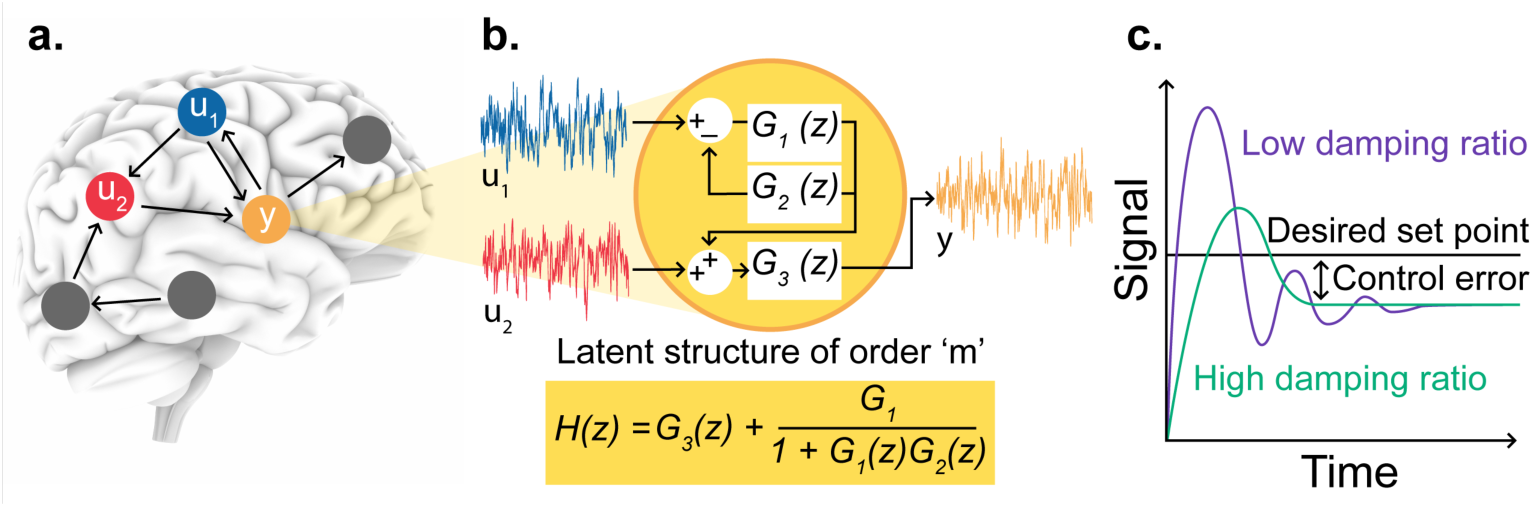
Overview of the proposed framework for quantifying control circuit regulation in the human brain. **a**. The first step involves obtaining the circuit architecture for selected nodes by establishing inputs to each node via causal inference, thus forming a directed network. **b**. The second step identifies circuit functionality using system identification to obtain a linear state-space model for each node. The directed network representation allows identifying each node’s transfer function as a multi-input single-output (MISO) system. For example, the MISO model for node y uses inputs from nodes u_1_ and u_2_. **c**. The final step involves probing the system using simulated inputs to quantify sensitivity to perturbations and measuring how well regulated or dysregulated the circuit is (control error). Systems with a low damping ratio are prone to stronger oscillatory behavior and thus more sensitive to perturbations than systems with high damping.

## Details of the proposed framework

### Functional MRI Data

We applied the methods described below to 5 hours of resting-state data for 10 subjects (mean age 29.1 ± 3.3 years, five women), collected over 10 subsequent days (30-minute resting-state scan each day) acquired on a Siemens TRIO 3T MRI scanner (Midnight Scanning Club, MSC^18^). Informed consent was obtained from all participants^18^. The study was approved by the Washington University School of Medicine Human Studies Committee and Institutional Review Board^18^. These included a total of four T1-weighted images (sagittal, 224 slices, 0.8-mm isotropic resolution, echo time (TE) = 3.74 ms, repetition time (TR) = 2400 ms, inversion time (TI) = 1000 ms, flip angle = 8°), four T2-weighted images (sagittal, 224 slices, 0.8-mm isotropic resolution, TE = 479 ms, TR = 3200 ms), four magnetic resonance angiographies (transverse, 0.6 × 0.6 × 1.0 mm, 44 slices, TR = 25 ms, TE = 3.34 ms) and eight magnetic resonance venographies, including four in coronal and four in sagittal orientations (sagittal: 0.8 × 0.8 × 2.0 mm thickness, 120 slices, TR = 27 ms, TE = 7.05 ms; coronal: 0.7 × 0.7 × 2.5 mm thickness, 128 slices, TR = 28 ms, TE = 7.18 ms). All functional imaging was performed using a gradient-echo echo-planar imaging (EPI) sequence (TR = 2.2 s, TE = 27 ms, flip angle = 90°, voxel size = 4 mm × 4 mm × 4 mm, 36 slices).

The data were spatially preprocessed in CONN toolbox^19^ by functional realignment and unwarping, slice-timing correction, segmentation, and Montreal Neurological Institute (MNI) normalization. The data were temporally preprocessed by physiological noise correction using CompCor^20^ (regressing five components of white matter and cerebrospinal fluid), regressing six motion parameters and bandpass filtering (0.008–0.1 Hz), followed by spatial smoothing with a 4-mm full width at half maximum Gaussian kernel.

### Identifying the Circuit Architecture

#### Selecting nodes

A node is the smallest entity of a circuit, representing a group of voxels that share similar temporal dynamics, with a neurobiological meaning and interpretability. “Node” is a term common within network science and graph theory, and it is commonly used in the functional connectivity literature interchangeably with “region of interest.” In our framework, a node acts as a brain region being controlled, a region exerting control, or both. In task-based designs, nodes are inferred via general linear modeling (GLM), whereas resting-state analyses typically select nodes using independent component analysis (ICA), with constraints on spatial properties (e.g., nodes may or may not overlap) and extent (whole-brain sampling vs. targeted sampling). Hypothesis-neutral (i.e., data-driven) methods include spatial ICA (sICA) for identifying spatially independent components, probabilistic functional modes^21^ (PROFUMO) for identifying modes of coherent activity, bottom-up parcellation schemes such as k-means clustering or spatially constrained hierarchical clustering^22^ for organizing smaller units into a region, and top-down parcellation using instantaneous connectivity^23^. However, for testing hypothesis-driven brain circuits^11^ or studying clinical populations with well-characterized regions of interest^13,14^ across the literature on brain disorders, predefined anatomical or functional nodes can generate comparable quantitative inferences. We used the Yeo parcellation^24^ to define cortical nodes and combined them with subcortical nodes obtained using the Harvard-Oxford atlas. Using predefined nodes can lead to the mixing of blood oxygen level–dependent (BOLD) time-series because of variability in brain structure across individuals or averaging of voxels contributing to cognitive process-induced activity with unrelated activity within a node. Therefore, we used the first eigen-variate (the highest variance-contributing component in the orthogonal decomposition of all time-series within a node) instead of an averaged time-series for a node.

#### Determining inputs

Identifying causal relationships and quantifying their strength from observed time-series in large-scale complex systems differs notably from undirected network reconstructions. With a known time-course for each node, interaction among nodes can be established by the presence of statistically significant correlation or by information-theoretic measures in the presence of nonlinear effects. Yet these fail to provide directionality or to specify direct versus transitive effects (i.e., the relationship *X* → *Y* → *Z* is incorrectly estimated as a triangular motif/network^25^). Several techniques to circumvent this statistical redundancy during network reconstruction exist; namely, partial correlation coefficient (pairwise correlation between nodes of interest after regressing out other nodes from both the nodes), conditional mutual information (link between two nodes is absent if mutual information between them, given other nodes, is statistically insignificant), and calculating probabilistic graphical models (zero elements of inverse covariance matrix, 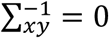, of a multivariate Gaussian distribution indicate an absent edge between nodes *x* and *y*). Identifying inputs requires capturing both temporal precedence and control. Lag-based methods (Granger causality, transfer entropy) using temporal precedence as the governing principle has been shown to perform poorly with fMRI data^26^, whereas methods based on statistical properties of the observed time-series (Patel’s Tau, Bayes Net methods) perform comparatively better^26^. Establishing causality using BOLD signal is notoriously difficult because of the slow hemodynamic response function, which low-pass filters neuron-driven dynamics. Further, the hemodynamic response varies across subjects and brain regions within a subject, interfering with the causal inference. Finally, the fMRI data exhibit a low signal-to-noise ratio^27^, which is typically addressed with preprocessing and denoising. However, these very steps used to amplify the signal can significantly distort its dynamics, and therefore causal inferences drawn from the dynamics.

Given the limitations of fMRI data, the most widely used and validated method to establish causality using fMRI has been DCM^28^. DCM uses a generative framework to model the hemodynamic response with hidden physiological states and a nonlinear observer function. DCM is causal in the control-theoretic sense, because it explains how the dynamics in one neural population causes the dynamics in another, as well as modulation of this interaction due to endogenous activity or experimental manipulation. Bayesian inversion provides posteriors for model parameters along with log model evidence to compare different DCMs with varying directed links among nodes. Recently, spectral DCM^17,29^ (spDCM) was introduced for modeling resting-state fMRI, which can invert large-scale graphs of the order of 36 to 64 nodes compared with fewer than 10 nodes with stochastic DCM^30^. Stochastic DCM inverts data in the time domain and requires estimating both time-variant hidden states and time-invariant model parameters, posing a computationally demanding inverse problem. Comparatively, spDCM inverts a deterministic model, requiring estimation of only time-invariant model parameters that explain the cross-spectra of observed data. The inversion efficiency is increased further by using functional connectivity modes to place prior constraints on undirected graph structure, followed by obtaining sparse representation using Bayesian model reduction of a fully connected graph^29^.

For the sample dataset, we identified inputs using spDCM for our 31 nodes for each subject’s 10 sessions. We then estimated the group parametric empirical Bayes (PEB) model using each subject’s data (10 sessions) and applied Bayesian model reduction to prune away parameters that did not contribute to model evidence and obtained a subject-specific effective connectivity matrix (**Fig. 2b**). In general, we found that feedback connections appeared to have stronger coupling than feedforward connections. We demonstrated that effective connectivity differs across each subject (**Fig. 2c**), which suggests that the feedback/feedforward control exerted on each node is individual-specific and must be inferred as such.

**Fig. 2:**
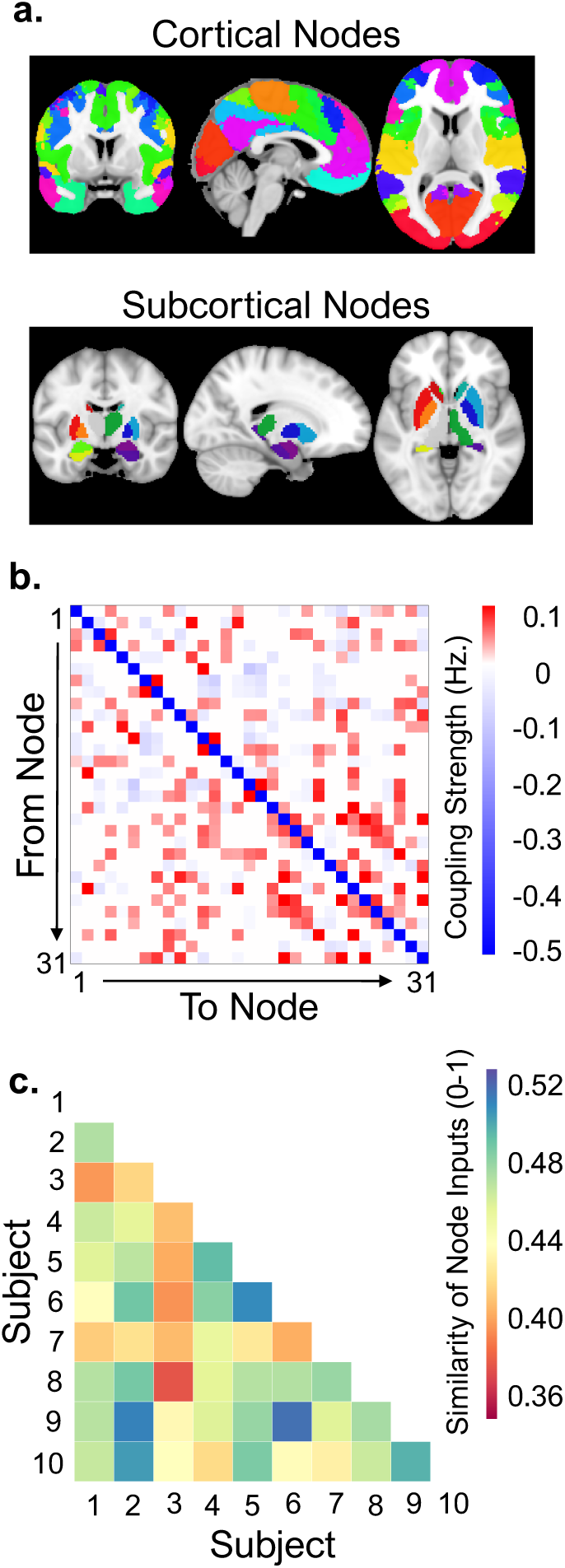
Large-scale resting-state dynamic causal modeling provides directed networks for obtaining the “input-output/cause-and-effect” relationship among nodes. **a**. Masks for cortical and subcortical nodes. **b**. Effective connectivity matrix for a representative subject, generated using spectral dynamic causal modeling, providing directed graphs. A circuit in the directed graph is simply a closed path formed by traversing the graph such that the first and the last node are the same, with no other node repeating during the traversal. **c**. Effective connectivity varies significantly across subjects; hence, for a given node, there is wide variation in inputs driving the node across subjects. The coupling similarity is calculated using the Dice coefficient on any two effective connectivity matrices.

### Identifying the Circuit Functionality

#### Generating transfer functions for each node

Establishing causality renders a multi-input single-output system, where a node receives input from other brain regions, with the measured time-series of the node as its response. We approximate the model for a given node using a discrete-time, linear time-invariant, stochastic system of model order *m* using

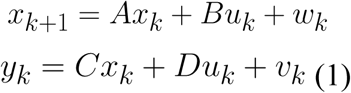

where *y*_*k*_ ∈ *R* is the measured BOLD signal of the node, *u*_*k*_ ∈ *R*^*n*^ corresponds to the input from other brain regions/nodes that were determined previously, and *x*_*k*_ ∈ *R*^*m*^, *w*_*k*_ ∈ *R*^*m*^, and *v*_*k*_ ∈ *R* represent the hidden node’s state, state noise (process noise), and measurement noise, respectively. *A* ∈ *R*^*m×m*^ is referred to as system matrix, *B* ∈ *R*^*m×n*^ as input matrix, *C* ∈ *R*^1*×m*^ as output matrix, and *D* ∈ *R*^1*×n*^ as feedthrough matrix. For an observable system, state estimation follows from

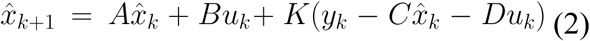

where *K* is the Kalman gain and

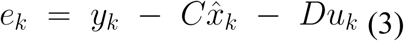

represents the white noise innovation process. Using equations (2) and (3), we can write the predictor form of equation (1) as

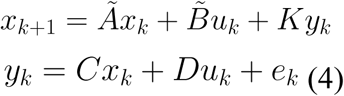

where *Ã* = (*A* – *KC*) and 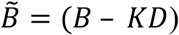. We seek to estimate free parameters such that given the input matrix, *u*_*k*_, and the initial state estimate,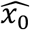, the model produces output 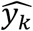 that approximates *y*_*k*_ with good accuracy. By iterating (4), we can write

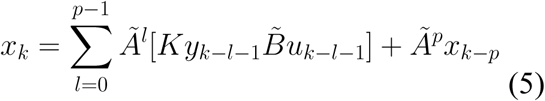

where *p* indicates the past horizon. We now define a finite future stacked vector of outputs with the future horizon, *f*, as 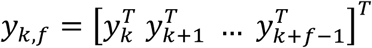. Finite future stacked vector of inputs, *u*_*k, f*_, and innovations, *e*_*k, f*_, follow the same definition. We can rewrite equation (4), using equation (5) and finite stacked vectors, as

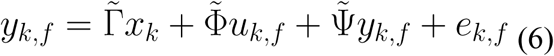

where

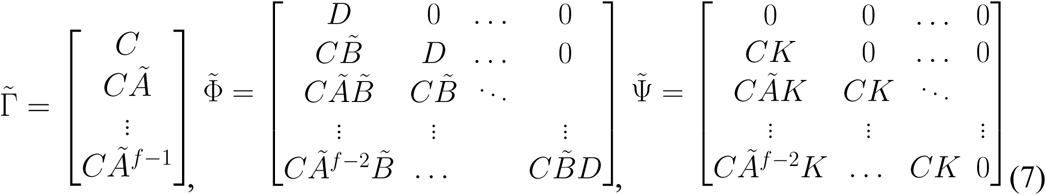

Importantly, note that 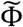 and 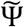 contain the predictor Markov parameters, 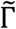 is the extended observability matrix, and choosing a large *p* cause *Ã*^*p*^ ≃ 0 with *x*_*k*_ represented by a linear combination of past inputs and outputs. Due to presence of feedback, *e*_*k,f*_ is correlated with *u*_*k,f*_ and hence we use the procedure specified by Jansson^31^ to pre-estimate a high-order ARX model and obtain unstructured estimates of the Markov parameters to form estimates 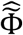 and 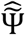. Using the estimates 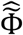 and 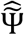, we can obtain

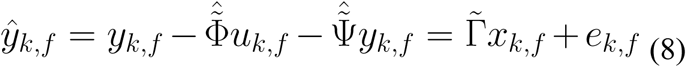

where the term *e*_*k,f*_ is uncorrelated with *u*_*k,f*_. The coefficient 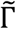 and unknown coefficients of past inputs and outputs of *x*_*k*_ (equation (5) with *Ã*^*p*^ ≃ 0) in equation (8) can then be estimated using least squares followed by singular value decomposition.

The model order, *m*, defines the model complexity and is a parameter of choice in the identification procedure. The choice of *m* can be based on either tuning *m* to obtain Markov parameter estimates that provide good prediction skill, as quantified by the correlation between the predicted 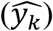 and the observed (*y*_*k*_) output, or by computing the number of dominant states, using Hankel singular values, that closely approximates the observed model response (*y*_*k*_). Hankel singular values provide a measure for energy of each state and are calculated as the square root of the eigenvalues for the product of the controllability and the observability Gramians.

#### Validating a putative circuit by comparing predicted trajectories against independent data

Estimated state-space models, using training data for each node across all subjects, showed high accuracy in approximating the fMRI output for the test data (**Fig. 3a**). Further, the dynamics were well approximated by low-complexity models (**Fig. 3b**). Increasing the number of data points in the system identification procedure improves the prediction skill and reduces uncertainty in the obtained parameter estimates (**Fig. 3c, 3d**). The state-space model (**Eq. 4**) of a node essentially represents coupled difference equation (or differential equation for continuous time), as an input-output transfer function expressed as *C*(*zI* − *A*)^−1^*B* + *D*. Therefore, the input-output transfer function represents a node as a computational unit built using three building blocks (**Fig. 4**).

**Fig. 3:**
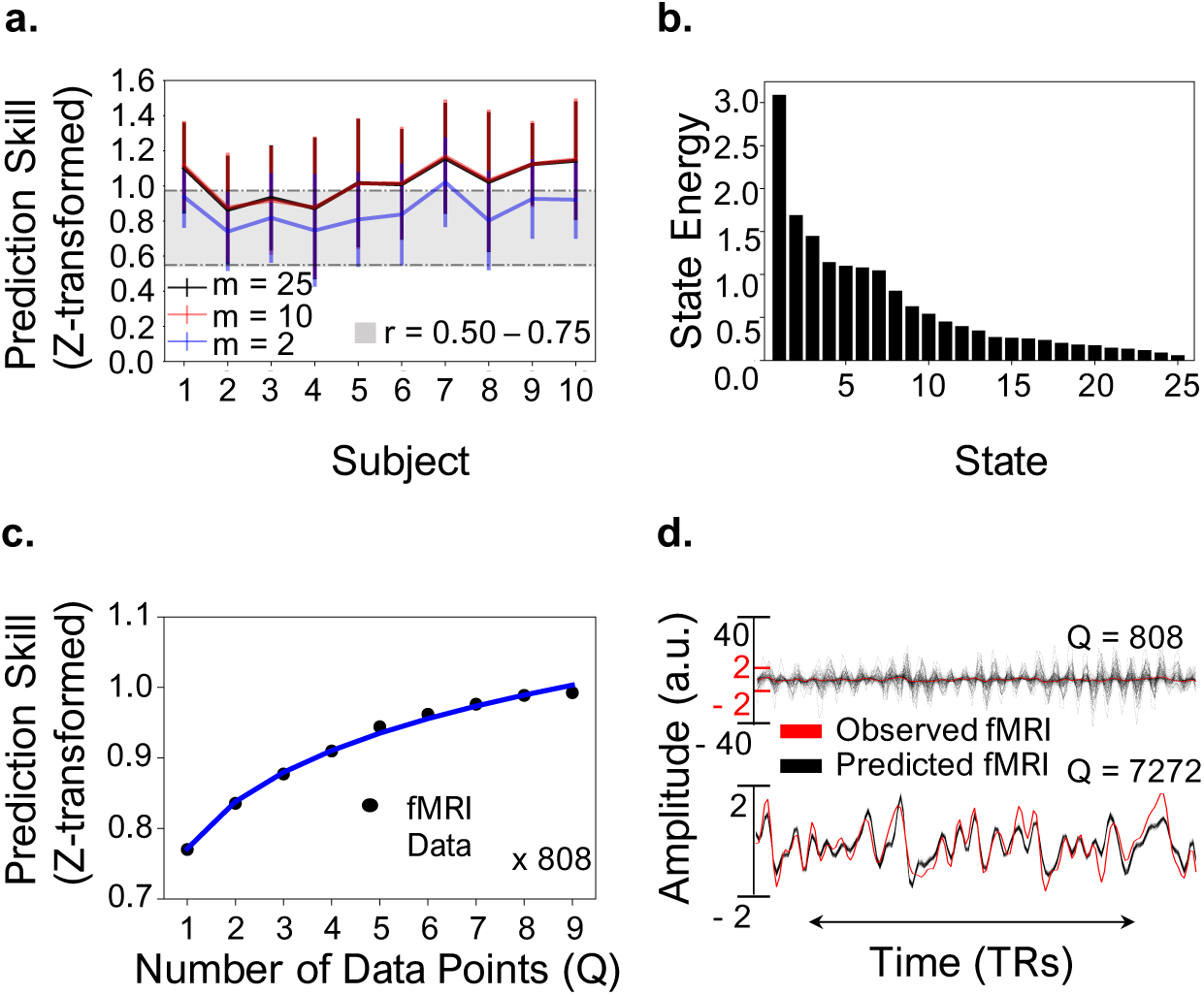
Node-level system identification provides a generative model for each node (brain region). **a**. Fisher Z-transformed correlation coefficient between the observed fMRI output, *y*_*k*_, and the predicted output,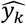(prediction skill), for the 10th scan for each node across all subjects at various model orders (*m*). System identification for estimating each node’s state-space model was performed using data from nine subsequent fMRI scans, and data from the 10th scan were used for testing (out-of-sample testing). Error bars indicate standard deviation across all 31 nodes. **b**. Hankel singular value decomposition for a node, for a representative subject, showing each state’s contribution in shaping the input-output behavior. Low-order models govern the input-output behavior and low-energy states can be discarded for obtaining simpler models, as evident also from a. (m = 10 vs. m = 25). **c**. Prediction skill, on 10th-day test data, as a function of the number of data points. Each point represents mean across all nodes and subjects; **d**. Predicted output,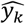 (black), for 100 Monte Carlo simulations using perturbations to uncertain model parameters. As the number of data points/time-series samples increase (from Q = 808 to Q = 7272) for system identification, model uncertainty decreases.

**Fig. 4:**
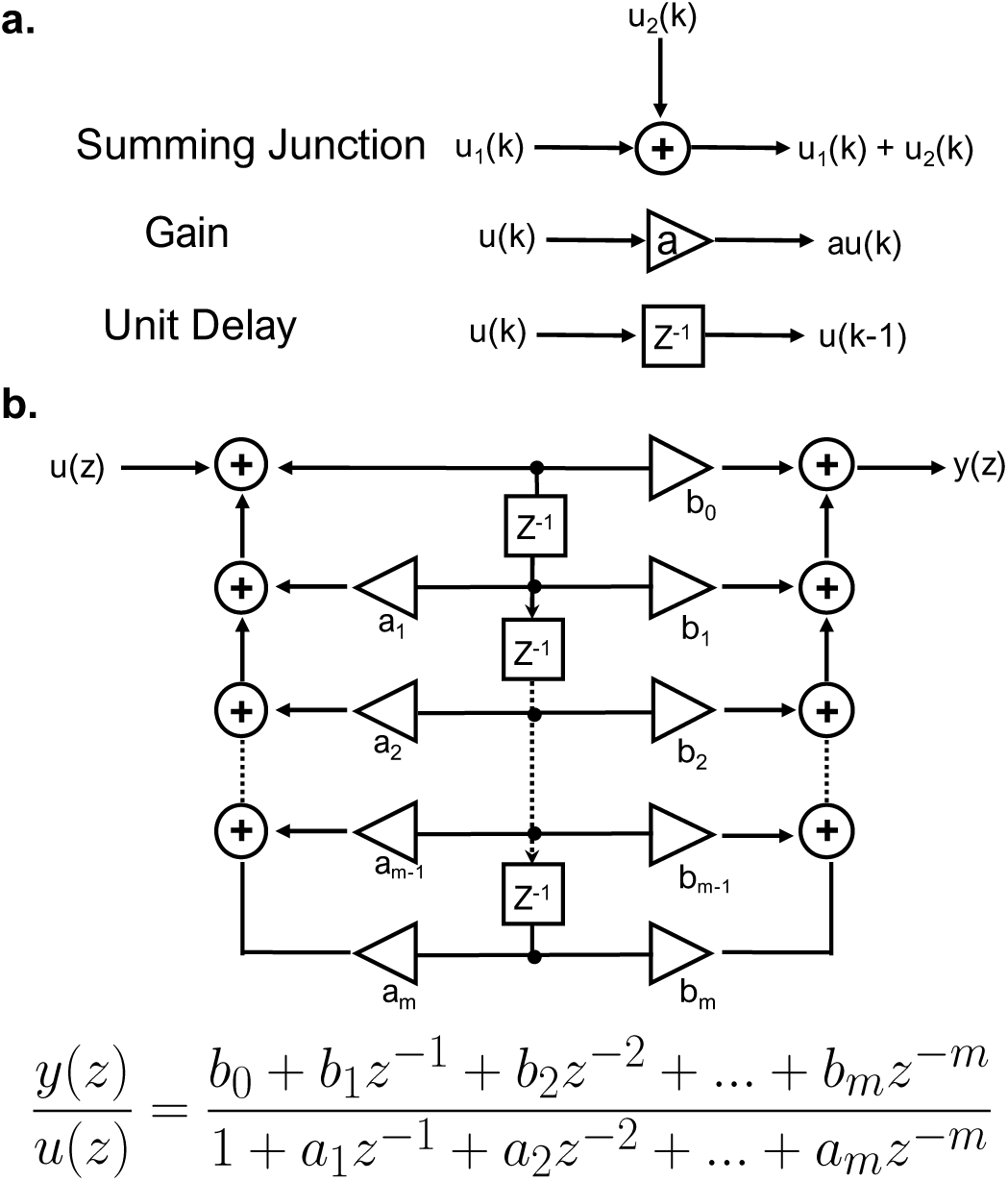
Implementing a synthetic circuit to mimic a node (brain region). **a**. Basic building blocks for realization, using hardware, of a transfer function or computation carried out by a node. The computational algorithm of a discrete linear-time-invariant system can be represented using the summing junction, gain, and unit delay/memory/storage element. **b**. Block diagram shows that any given *m*th-order transfer function can be realized using the basic building blocks. If the number of delay elements in a realization using hardware is equal to the order of the difference equation, the realization structure is called a canonical structure.

### Probing the System

#### Calculating a circuit’s sensitivity to external perturbation

Neural information processing is shaped by oscillatory dynamics that support cognitive control by modulating excitability in neuronal populations^32,33^. The oscillatory dynamics could amplify, sustain, or dampen a given input for efficient cognitive processing, with abnormal dynamics inducing psychopathology. For example, dysfunctional neural synchrony due to deficits in γ-aminobutyric acid (GABA) neurotransmission has been associated with cognitive dysfunction in schizophrenia^34^. The dynamic states associated with each node in a circuit are regulated by its neural activity, local neurotransmitters, and the balance between inhibitory-excitatory control inputs^35^, along with coupling strength and timing delay. Probing the learned dynamics of a node through state-space identification using fMRI measurements combines the effect of regional neural activity, local neuromodulation, and the excitatory or inhibitory control inputs to the node. With available state-space models for each node (**Eq. 1-8**), we can estimate the regulated behavior (both self-regulation and control exerted by feedback/feedforward control inputs) of each node using information about the eigenvalues of the system matrix, A. Location of the eigenvalues (poles of a transfer function are the eigenvalues of the system matrix), in the s-plane (Laplace domain) for continuous time or z-plane for discrete-time, provides the damping ratio and the natural frequency of a given system (**Fig. 5**).

**Fig. 5:**
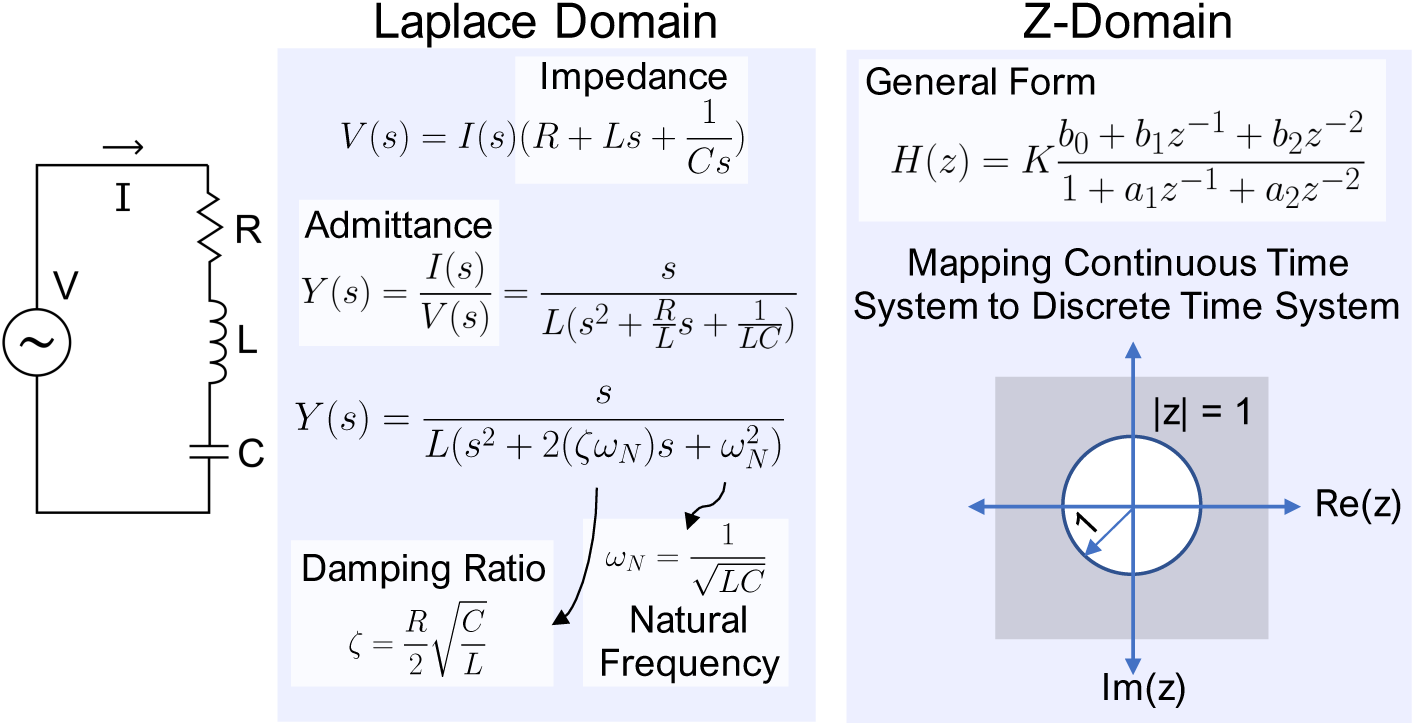
Location of the poles provides an estimate of the damping ratio and the natural frequency of a system. A series resistor-inductor-capacitor (RLC) circuit is a damped harmonic oscillator and described by a second-order differential equation or a second-order transfer function (admittance) in Laplace domain (continuous time, s-domain) or Z-domain (discrete time, as for fMRI measurements). Z-transform maps a continuous-time system to a discrete-time system under the transformation *z* = *e*^*sT*^ for a sampling time period T. The imaginary axis of the s-plane maps on the unit circle |z| = 1, the left half of the s-plane maps inside the unit circle |z| = 1, and the right half of the s-plane maps outside the unit circle in the Z-domain. The poles of the transfer function, computed by finding roots of the characteristic equation, determines the oscillatory behavior of the system. For example, for a second-order system, conjugate poles on imaginary axis (s-plane) or on the unit circle (Z-domain) lead to an undamped oscillator. In general, for a discrete-time system with pole location z and sampling time T, the damping ratio is given by *ζ* = −*cos*(∠*ln*(*z*)) and natural frequency is given by 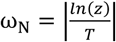.

Tracking damping ratio and natural frequency of nodes longitudinally for a subject provides a way to understand trajectories of clinical onset and relapse^36^, including changes to circuit dynamics due to pharmacological intervention^37^. For a given node, the damping ratio and frequency ratio (ratio of the input frequency versus natural frequency) induce phase shift in an incoming input from another brain region, as shown in **Fig. 6A**. This phase shift, in turn, affects functional connectivity, measured by pairwise Pearson correlation between nodes. We show how the functional connectivity between a pink-noise input signal and its corresponding phase-shifted output varies for a second-order resistor-inductor-capacitor (RLC) circuit (**Fig. 6B**) at different damping ratios and natural frequencies. We further show the distribution of functional connectivity between inputs (having pink-noise spectra) to a node and its corresponding output for a sample subject (**Fig. 6D**), illustrating how node dynamics explain synchrony or asynchrony with its input nodes. Changes in synchrony arising because of self-regulation, or rerouting of connections as quantified by causality analysis, or neuromodulatory effects of a pharmacological intervention are easily quantified by applying simulated inputs to a node’s generative model built across time and in response to treatment.

**Fig. 6:**
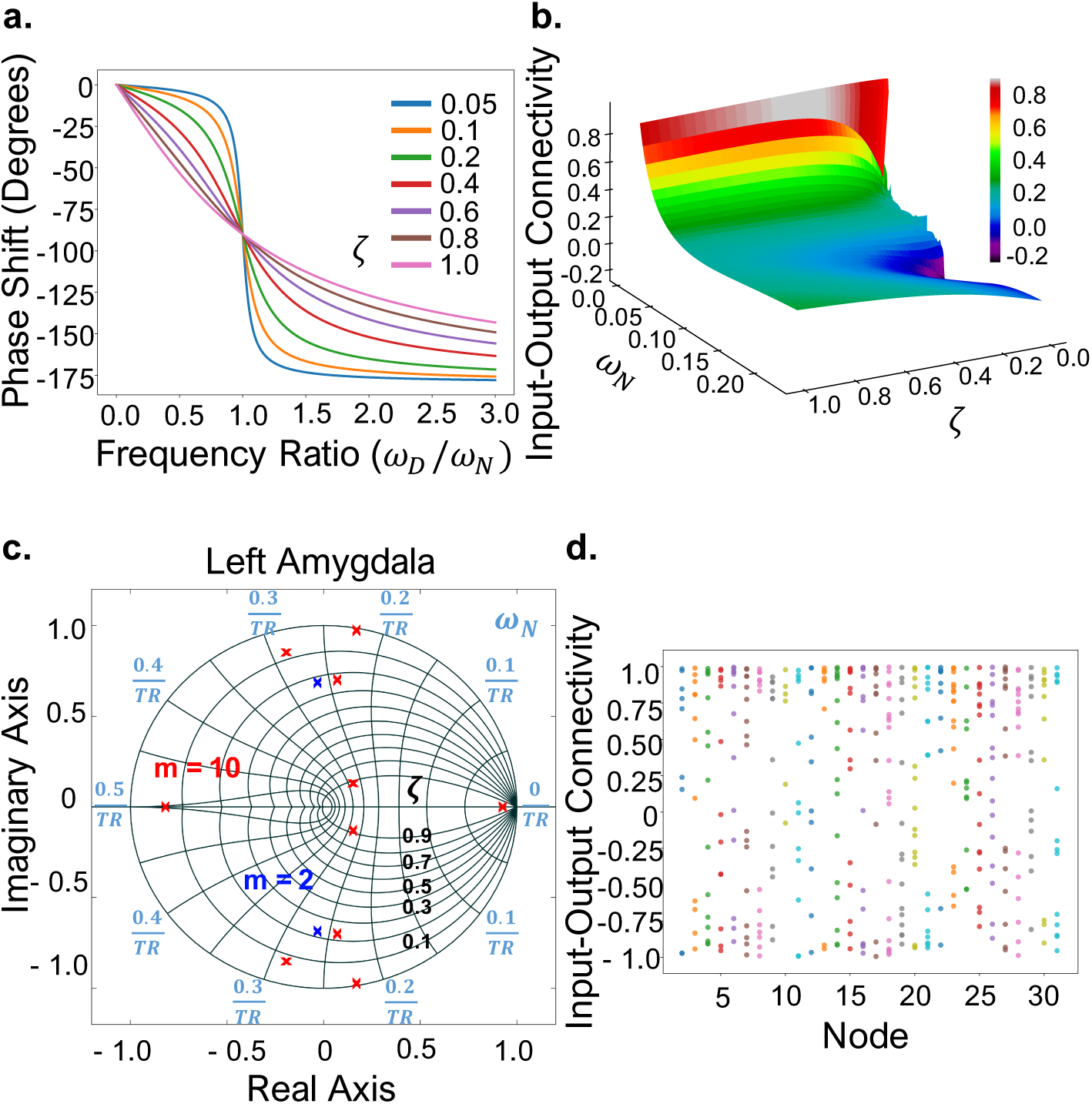
Damping ratio and the natural frequency modulate connectivity among brain regions. **a**. When subjected to a sinusoid input, a linear-time invariant system can induce a change in the amplitude or the phase at the output while maintaining the same output frequency as the input. The phase shift is dependent on the damping ratio and the frequency ratio (ratio of driving input frequency, *ω*_*N*_, and the natural frequency, *ω*_*D*_), as shown for the second-order series resistor-inductor-capacitor (RLC) circuit. **b**. Effect of phase shift by quantifying Pearson correlation (connectivity) between a pink-noise input signal and its corresponding output for the series RLC circuit, at varying damping ratio and natural frequencies. **c**. The damping ratio and natural frequency of left amygdala for a representative subject at two different model orders, *m* = 2 (blue) and *m* = 10 (red). Lower complexity models enable comparison with standard engineering control systems; however, they suffer from low prediction accuracy (Fig. 3a). **d**. Simulating each node’s response to pink-noise inputs (all inputs to a node are distinct and randomly generated) followed by calculation of Pearson correlation (connectivity) between each input and the output of the node, for a representative subject. The input-output pairs represent nodes that can show synchrony, anti-synchrony, or no synchrony characterized by the input-output transfer function.

Brain circuits must ideally maintain stability while also being responsive to relevant stimuli^35^. For an intuitive understanding, we demonstrate how the damping ratio and frequency ratio affect magnitude gain and hence sensitivity to inputs in a second-order resistor-inductor-capacitor (RLC) circuit (**Fig. 7b**). We extend the analysis to observing the sensitivity of brain regions to input perturbations of varying frequency content (healthy resting-state fMRI signals follow pink-noise frequency spectra). Inputs with frequency spectra other than pink noise simulate the effect of a perturbation by environmental noise, with **Fig. 7c** showing more sensitive regions that have an abnormally high magnitude gain at non-pink-noise inputs (e.g., regions 10 and 27 for subject 1, and regions 6, 18, 25, and 29 for subject 10 in **Fig. 7c**).

**Fig. 7:**
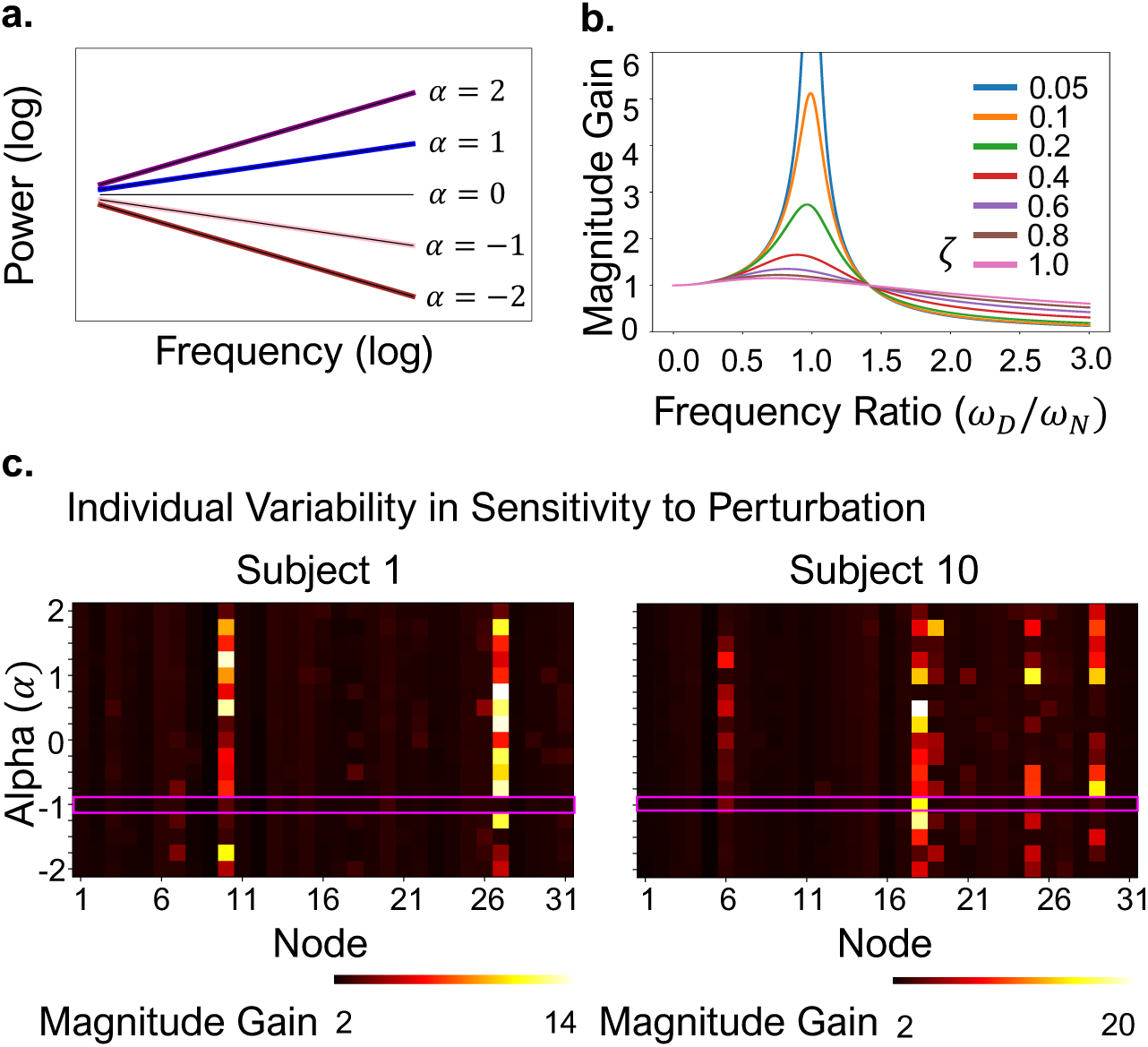
Measuring sensitivity to external perturbations using magnitude gain. **a**. Spectra for various colored (power law, *f*^*α*^) noise. **b**. Effect of damping ratio and the frequency ratio on magnitude gain/amplitude modulation of the input sinusoid for the series resistor-inductor-capacitor (RLC) circuit. **c**. Magnitude gain for colored-noise inputs (healthy brain exhibits pink-noise spectra) for testing sensitivity to external perturbations. The representative subjects show abnormally high magnitude gain at non-pink-noise inputs for certain nodes, with node 18 for subject 10 showing abnormal behavior even for a pink-noise input, indicating a potential failure/susceptible point. Sensitivity can be situational and, thus, can be used as a parameter of interest for comparison across the spectrum of psychiatric illness (e.g., comparing sensory gating in schizophrenia spectrum on a continuum from severe, chronic schizophrenia to attenuated schizophrenia^40^).

#### Calculating control error as a quantifiable measure of regulation/dysregulation of the circuit

Feedforward input to a system pushes the system in a predetermined direction, without knowing the error signal (the difference between the desired and the actual state). In contrast, the feedback control actuates the system based on the error signal in a direction that reduces the error at each consecutive time-step. A commonly used performance metric for the design and optimization of controllers (e.g., a proportional–integral–derivative controller) in engineering control systems is the steady-state error (the error of a system as the time approaches infinity). The control enacted on each node/brain region combines both the feedback and feedforward control inputs, and we can calculate the steady-state error or efficacy of the enacted control using the final value theorem with a step input. For a final steady-state value *F*, the control error (CE) is calculated as *CE* = |1 − *F*| (**Fig. 8a**). We demonstrate that CE varies across regions for each subject (**Fig. 8b**). For a single subject, tracking CE longitudinally will provide an assessment of coupling efficiency between brain regions in a circuit for enacting control. A systematic increase in steady-state error longitudinally would indicate circuit-level dysregulation.

**Fig. 8:**
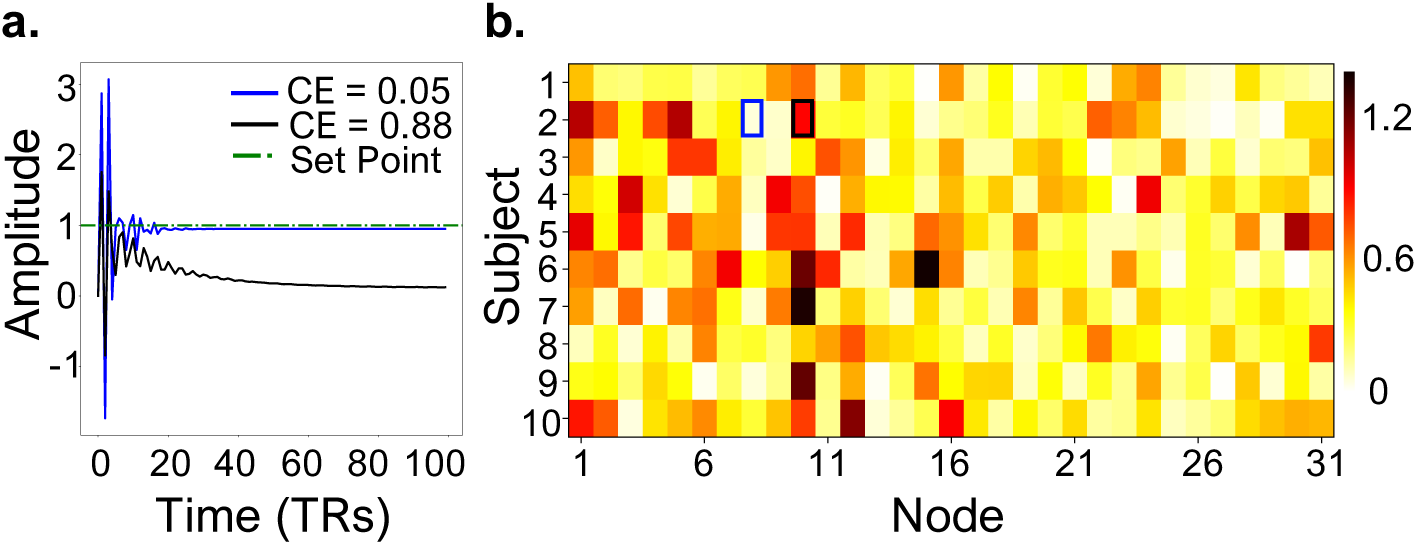
Quantifying regulation/effectiveness of endogenous control using control error. **a**. Steady-state response of two nodes, for a representative subject, showing varying amounts of deviation from the desired set point. The closer the final steady-state value is to 1, the lower the control error, and the better the circuit regulation. **b**. Control error (CE) calculated for each node for a unit-step input using the final value theorem, showing wide variance among nodes and across subjects. The blue and black boxes for subject 2 correspond to the control error demonstrated in a.

## Conclusion

Neuroimaging analytic tools are conceptually and mathematically optimized for quantifying activation and connectivity, rather than circuits and their regulation. Here, we provide a proof of concept for how control systems engineering techniques may be adapted to fMRI, to extend our neuroimaging capabilities in quantifying circuit dynamics. In our simple model, we demonstrate how we can quantify two key features of direct relevance to understanding onset and relapse in psychiatric disorders: a circuit’s sensitivity to perturbation and its degree of dysregulation. This framework may be expanded to include more complex architectures (e.g., linked positive and negative feedback loops) and additional functions (e.g., gating, gain and lag), as required to model specific circuits and/or disorders^38^. One important consequence of generative models is that they permit simulations, and therefore prediction, of future trajectories under various conditions. Thus, control circuit models go beyond descriptive or conceptual utility, permitting us to more rigorously validate (and falsify) models than current statistical practice^39^.

While we present these techniques here in the context of fMRI, the system identification process is streamlined and improved with the superior temporal resolution and dynamic fidelity of time-series made possible by magnetoencephalography (MEG), electroencephalography (EEG), or local field potentials (LFP). We started with fMRI for two reasons. First, psychiatry-specific control circuits typically rely critically upon both cortical and subcortical components. The reliance on high-fidelity subcortical time-series introduces technical challenges for cortical modalities such M/EEG, and the nonhomologous structure of the neocortex across species is a different challenge for invasive modalities such as LFP. Second, fMRI is currently the dominant experimental tool in human neuroscience, and psychiatric disorders are still generally understood as human disorders. Thus, although fMRI is the hardest use-case for quantifying control circuit regulation, it is also the modality in which achieving this goal can have the greatest clinical impact.

## Acknowledgments

The research described in this paper was funded by the Baszucki Brain Research Fund, and the National Institute of Drug Abuse (SBIR Phase 1 & Phase 2 Grant 1R44 DA043277-01). The authors gratefully acknowledge the assistance provided by Adeel Razi (Monash University, Australia) for discussions on dynamic causal modeling.

## Authorship Contribution Statement

L.R.M.-P., R.K., and H.H.S. conceptualized the study. R.K. developed the proposed framework and analyzed the data. R.K., L.R.M.-P., and H.H.S. wrote and revised the manuscript. L.R.M.-P. and H.H.S. acquired the funding.

## Data Availability

The fMRI data used in this work were obtained from OpenNEURO and are available from https://openneuro.org/datasets/ds000224/versions/1.0.3.

## Code Availability

All source code is available from the corresponding author upon reasonable request.

